# Uncovering *Fusarium* species diversity, distribution and pathogenicity associated with cotton-corn rotation fields in recently expanded cotton-growing areas of the Northern High Plains of Texas

**DOI:** 10.1101/2025.07.02.662791

**Authors:** Ihsanul Khaliq, Nolan R. Anderson

## Abstract

The recent expansion of cotton acreage in the Northern High Plains of Texas has raised concerns about the occurrence of a diverse *Fusarium* community, particularly *Fusarium oxysporum* f. sp. *vasinfectum* race 4. We investigated the diversity, distribution, and pathogenicity of *Fusarium* species in soil samples collected from 26 cotton-corn rotation fields across 10 counties. A total of nine *Fusarium* species were isolated: *F. incarnatum*, *F. equiseti*, *F. solani*, *F. verticillioides*, *F. proliferatum*, *F. clavum*, *F. oxysporum*, *F. flocciferum*, and one unidentified *Fusarium* sp. The highest species diversity (five species) was observed in Sherman County. *Fusarium solani* was the most frequently isolated species (39%) across all counties, followed by *F. equiseti* (22%), while *F. flocciferum* was the least frequently isolated species (2%). Greenhouse pathogenicity assays showed *F. incarnatum* caused significantly higher disease severity relative to non-inoculated control on aerial parts of cotton (*P*=0.0137), while *Fusarium flocciferum*, *F. proliferatum,* and *F. clavum* caused significantly higher disease severity relative to non-inoculated control on roots (*P=*0.01, *P=*0.01, *P=*0.02, respectively). On corn, differences in disease severity were not statistically significant (*P*>0.05). *Fusarium* isolates were recovered from both symptomatic and asymptomatic roots following pathogenicity trials, suggesting endophytic colonization. This study revealed a complex *Fusarium* community capable of cross-infecting cotton and corn, with implications for regional disease management. The absence of *Fusarium oxysporum* f. sp. *vasinfectum* race 4 indicates it may not yet have been introduced into the Northern High Plains; however, continued surveillance is warranted given its detection in New Mexico and adjacent counties in Texas.

---

The Northern High Plains are a mixture of rangeland and cropland, comprising the 22 northernmost counties in Texas (Zivkovic and Hudson 2012). The region faces severe water scarcity, with an annual mean precipitation of about 558 mm–well below the US national average of 965 mm per year (BestPlaces 2025; NOAA 2021).

Additionally, the region relies heavily on water drawn from the Ogallala Aquifer to produce irrigated crops. However, current water use is unsustainable, with annual extraction exceeding annual recharge (Thayer and McCarl 2018). Approximately 7 million beef cattle are raised annually in beef cattle feedyards in the Texas High Plains (TCFA 2015). As of March 2025, there were 2.35 million head of cattle and calves on feed in the Northern High Plains, accounting for 89 percent of the state’s total (USDA National Agricultural Statistics Service 2025). Corn remains the primary feed crop, with more than 50 percent of the corn grown in Texas coming from the High Plains region (National IPM Database 2001; Xue et al. 2017). Although corn is the major feed crop, there has been significant interest in growing cotton as a rotational crop due to water constraints and the high fertilizer costs associated with corn production (Colaizzi et al. 2009).

Cotton (*Gossypium* species) is the most important fiber crop for the textile industry and a significant oilseed crop globally. The United States is the world’s fourth-largest cotton producing country (USDA Foreign Agricultural Service 2024), with cultivation spanning 17 southern states. Texas is the leading producer, accounting for approximately 40% of the U.S. cotton production in recent years (USDA Economic Research Service 2022). Although cotton was introduced to the Northern High Plains in 1968, acreage remained below 20,000 ha until 1998, but has expanded substantially since 2004—including a 126% increase between 2013 and 2019 alone—raising concerns about diseases that may affect cotton production (Ledbetter 2020; Marek and Bordovsky 2006).

Key fungal genera implicated in cotton diseases worldwide and in the U.S. Cotton Belt include *Fusarium*, *Verticillium*, *Rhizoctonia*, and *Pythium*. Among these, *Fusarium*, *Rhizoctonia*, and *Pythium* species are primarily associated with the seedling disease complex (Howell 2002; Watkins 1981), while *Fusarium* species—alone or in combination with *Alternaria*, *Aspergillus* species, or bacterial pathogens such as *Pantoea agglomerans* and *Serratia marcescens*—are frequently linked to seed rot (Glover et al. 2020; Medrano et al. 2007; Minton and Garber 1983). Diseases caused by oomycetes and splash dispersed pathogens are not considered a major concern in the Northern High Plains due to the dry climate and severe water scarcity.

Fusarium wilt, caused by *Fusarium oxysporum* f.sp. *vasinfectum* (FOV), is of particular concern in the Northern High Plains because cooler temperatures during the early planting season favor its development (Zhang et al. 2020). Night temperatures in the Northern Panhandle are among the coolest in cotton growing regions of the U.S., with mean annual temperature of about 11°C (Kimmel Jr et al. 2016). FOV was first detected in the United States in 1892 (Atkinson 1892). Historically, FOV has been characterized into 8 different races based on variation in pathogenicity to various plant species, including *Gossypium* species and cultivars. Among these, race 4 (FOV4) is of particular concern in the U.S. due to its high virulence on both Upland (*G. hirsutum* L.) and Pima (*G. barbadense* L.) cotton, as well as its ability to persist in soil for years as chlamydospores (Hutmacher et al. 2013; Kim et al. 2005). Additionally, unlike other races, FOV4 can cause severe wilting in the absence of root knot nematode *Meloidogyne incognita.* FOV4 was first detected in the U.S. in 2001 in California (Kim et al. 2005) and more recently in 2017 in Texas (Halpern et al. 2018) and 2019 in New Mexico (Zhu et al. 2019). This highly aggressive race has not yet been reported in the Northern High Plains. However, given the proximity to New Mexico and adjacent counties in Texas where FOV4 has been detected (Cianchetta et al. 2015; Diaz et al. 2021), there is great concern about its potential introduction into the Northern High Plains. Therefore, a comprehensive survey is urgently needed to investigate the presence or absence of FOV4 in the region.

Several surveys have been conducted across the U.S. Cotton Belt for *Fusarium* species (Cianchetta et al. 2015; Diaz et al. 2021; Mao and Hu 2019; Zhu et al. 2021). However, these surveys did not include the Northern High Plains, leaving limited information on the diversity, distribution and relative severity of *Fusarium* species affecting cotton in this region. Moreover, these studies were primarily focused on investigating the presence or absence of FOV4 rather than determining overall *Fusarium* species diversity. Several *Fusarium* species are known to cause economically important diseases in cotton, including wilt, damping-off, boll rot, and root rot (Davis et al. 2006; Zaki et al. 2021). While *F. oxysporum* and *F. solani* are the most common species complexes associated with cotton, the relative severity and impact of other *Fusarium* species remain poorly understood (Mao and Hu 2019).

Crop rotation has been recognized as an effective practice to reduce nitrogen fertilizer use and enhance both grain yield and sustainability in agricultural systems (Shu et al. 2021). However, crop rotation can also inadvertently increase inoculum levels, thereby elevating the risk of crop diseases (Li et al. 2021). Previous studies have demonstrated that crop residue serves as a natural reservoir for *Fusarium* species and may even facilitate genetic exchange (Ward et al. 2008). For instance, in wheat corn cropping rotations, continuous wheat cultivation following corn has been shown to increase the incidence of *Fusarium* stalk rot and ear rot in maize, as well as *Fusarium* head blight in wheat (Xi et al. 2021; Zhang et al. 2016). In the Northern High Plains, where cotton and corn are commonly grown in rotation, investigating the pathogenicity of *Fusarium* species on both crops is crucial. Such studies will provide valuable insights into the pathogen’s persistence, cross-infectivity, and the associated epidemiological risks.

Greenhouse pathogenicity assays offer several advantages, including reduced variability resulting from uneven inoculum distribution and external biotic or abiotic factors. Most pathogenicity assays conducted in greenhouse conditions use root-dip inoculation method, where two-week-old seedlings’ roots are dipped in a spore suspension for two minutes and then transplanted into healthy soil (Kim et al. 2005). This method can predispose seedlings to stress during transplantation and does not reflect host infection in the field (Schoeneweiss 1975). Conversely, pathogenicity assays that involve directly planting seeds in *Fusarium*-infested soil or adding inoculum after seedling emergence better reflect infection under field conditions (Diaz et al. 2021).

To address these key gaps, the present study was conducted to: (1) conduct a comprehensive survey to determine the diversity and distribution of *Fusarium* species associated with cotton-corn rotation fields in the recently expanded cotton-growing areas of the Northern High Plains; (2) investigate the presence or absence of FOV4 in the region; and (3) evaluate the pathogenicity and severity of *Fusarium* species on cotton and corn under greenhouse conditions.

## Materials and Methods

### Sampling sites and sampling procedure

With the help of growers and county agents, a total of 26 cotton-corn rotation fields across 10 different counties in the Northern High Plains of Texas were sampled between late spring and late fall (May to October) of 2024. Specifically, six fields were sampled in Sherman County; four fields each in Moore and Deaf Smith counties; three fields in Hartley County; two fields each in Randall, Hansford, and Carson counties; and one field each in Potter, Dallam, and Hutchinson counties. Fields selection was based on growers or county agents’ willingness to participate in the study.

In each field, a total of 25 soil cores, each 0.2 m deep, were collected at 5 different locations, with at least 60 m space between sampling points (5 soil cores per sampling location within a field, 25 soil cores per field). The soil samples from each field were pooled and placed into cloth bags, kept on ice in an insulated box to protect them from high temperature and direct sunlight, and carried to the laboratory.

In the laboratory, the samples collected from each field were mixed thoroughly and air dried for a week to inhibit microbial activity and facilitate grinding and sieving (Sangalang et al. 1995). The air-dried soil sample for each field was poured into a Nasco-Asplin soil grinder, pulverized gently to break the clods, and passed through a 2 mm sieve. To prevent cross-contamination between samples, the excess dirt from grinder’s parts were removed with a brush, washed with water, and then cleaned thoroughly with 70% ethanol and paper towels.

### Diversity and distribution of *Fusarium* species in the Northern High Plains Isolation of single spore isolates from soil

To isolate *Fusarium* species from soil, a total of 20 cc of soil was added to 80 mL of reverse osmosis (RO) water in a small beaker (200 mL) placed on a magnetic stirrer and mixed vigorously to ensure a thorough mixing of the soil. From this soil suspension, a 1 mL aliquot was plated onto each of 5 Petri dishes of Peptone PCNB agar (PPA) medium (1000 mL distilled water, 15 g Peptone, 1 g KH_₂_PO_₄_, 0.5 g MgSO_₄_·7H_₂_O, 0.75 g PCNB, 20 g agar, pH adjusted to 5.5–6.5) (Nash and Snyder 1962). The process was repeated with an additional 20 cc of soil using another set of five Petri dishes, allowing us to assess 40 cc of soil for each field using a total of 10 Petri dishes. The Petri dishes were incubated for 3 days on PPA at room temperature (23 ± 2°C) with a 12-hour photoperiod. The colonies developed on PPA were subcultured onto full strength potato dextrose agar (PDA) medium by taking small mycelial plugs, observed under a Leica M205C stereoscope (Leica Camera AG, Wetzlar, Germany) at 160x magnification, from the colony margins.

After incubating for two weeks on PDA, 200 μL of RO water was added to each Petri dish, and conidia were dislodged by gently scrapping the mycelium with a pipette tip. A total of 150 μL of this conidial suspension was then transferred onto 30% water agar and incubated overnight. Single spore isolates were obtained by transferring single germinated conidia, observed under Nikon SM225 microscope (Nikon corp., Tokyo, Japan) at 180x magnification, onto fresh PDA plates.

### Genomic DNA extraction, PCR amplification, and sequencing

Isolates were divided into morphotypes based on their gross colony morphology (e.g., mycelial growth patterns), and their identities were obtained using the nuclear ribosomal internal transcribed spacer region (ITS rDNA) and the translation elongation factor 1-α (tef-1α) gene region (Table 1). DNA was extracted using Zymo Quick DNA Fungal/Bacterial Microprep kit (Zymo Research Corp; Irvine, CA). Amplicons were obtained using the following primer pairs: ITS1 (TCCGTAGGTGAACCTGCG) × ITS4 (TCCTCCGCTTATTGATATGC) for ITS rDNA (White et al. 1990); and EF-1 (5′-ATGGGT AAGGARGACAAGAC-3′) × EF-2 (5′-GGARGTACCA GTSATCATG-3′) for tef-1α gene region (O’Donnell et al. 1998). The PCR reaction mixture (25 μl reaction) contained 12.5 μl of Phire Green Hot Start II Master Mix (Thermo scientific), 1.25 μl of each primer (0. 5 μM), and 1.5 μl of genomic DNA. The PCR amplification program conditions were as follows: initial denaturation at 98°C for 90 s followed by 35 cycles at 98°C for 5 s, annealing at 58°C for 15 s and extension at 72°C for 15 s, followed by final extension for 4 minutes at 72°C.

**Table 1.** Isolate identity, collection date, county of origin, host crop history, and GenBank accession numbers for *Fusarium* species isolated in this study.

| Isolate | Organism | County | Previous crop | Current crop | Sampling date | GenBank Accession number |  |  |
| --- | --- | --- | --- | --- | --- | --- | --- | --- |
| | | | | | | tef-1α | ITS | RPB2 |
| F1 | *F. incarnatum* | Sharman | Corn | Cotton | 2024/08/22 | PV699158 |  |  |
| F10 | *F. incarnatum* | Moore | Corn | Cotton | 2024/08/21 |  | PV670408 |  |
| F2 | *F. equiseti* | Carson | Corn | Cotton | 2024/10/24 |  | PV670407 |  |
| F14 | *F. equiseti* | Carson | Corn | Cotton | 2024/10/24 | PV699144 |  |  |
| F15 | *F. equiseti* | Moore | Corn | Cotton | 2024/08/21 | PV699145 |  |  |
| F18 | *F. equiseti* | Hartley | Corn | Cotton | 2024/09/26 | PV699148 |  |  |
| F20 | *F. equiseti* | Sharman | Corn | Cotton | 2024/08/22 | PV699152 |  |  |
| F25 | *F. equiseti* | Hansford | Cotton | Wheat | 2024/05/01 | PV699157 |  |  |
| F38 | *F. equiseti* | Hutchinson | Fallow | Cotton | 2024/05/21 | PV717850 |  |  |
| F40 | *F. equiseti* | Sharman | Corn | Cotton | 2024/08/22 | PV717843 |  |  |
| F41 | *F. equiseti* | Sharman | Corn | Cotton | 2024/08/22 | PV717844 |  |  |
| F3 | *F. solani* | Deaf Smith | Fallow | Cotton | 2024/10/02 | PV699137 |  |  |
| F4 | *F. solani* | Deaf Smith | Corn | Cotton | 2024/10/02 | PV699138 |  |  |
| F7 | *F. solani* | Deaf Smith | Corn | Cotton | 2024/10/02 | PV699140 |  |  |
| F8 | *F. solani* | Moore | Corn | Cotton | 2024/08/21 | PV699141 |  |  |
| F9 | *F. solani* | Moore | Corn | Cotton | 2024/08/21 | PV699142 |  |  |
| F19 | *F. solani* | Carson | Corn | Cotton | 2024/10/24 | PV699149 |  |  |
| F21 | *F. solani* | Deaf Smith | Corn | Cotton | 2024/10/02 | PV699153 |  |  |
| F22 | *F. solani* | Carson | Corn | Cotton | 2024/10/24 | PV699154 |  |  |
| F23 | *F. solani* | Deaf Smith | Wheat | Cotton | 2024/10/02 | PV699155 |  |  |
| F30 | *F. solani* | Moore | Corn | Cotton | 2024/08/21 | PV717836 |  |  |
| F31 | *F. solani* | Hutchinson | Fallow | Cotton | 2024/05/21 | PV717837 |  |  |
| F32 | *F. solani* | Sharman | Corn | Cotton | 2024/08/22 | PV717838 |  |  |

Plant Disease
| Isolate | Organism | County | Previous | Current | Sampling | GenBank Accession number |  |  |
| --- | --- | --- | --- | --- | --- | --- | --- | --- |
| F33 | *F. solani* | Dallam | Corn | Cotton | 2024/09/26 | PV717839 |  |  |
| F35 | *F. solani* | Randall | Corn | Cotton | 2024/08/22 | PV717840 |  |  |
| F36 | *F. solani* | Hartley | Corn | Cotton | 2024/09/26 | PV717841 |  |  |
| F37 | *F. solani* | Carson | Corn | Cotton | 2024/10/24 | PV717842 |  |  |
| F5 | *F. verticillioides* | Sharman | Corn | Cotton | 2024/08/22 | PV699139 |  |  |
| F42 | *F. verticillioides* | Sharman | Corn | Cotton | 2024/08/22 | PV717845 |  |  |
| F12 | *F. proliferatum* | Randall | Corn/Sorghum | Cotton | 2024/08/22 |  | PV670409 |  |
| F16 | *F. proliferatum* | Randall | Corn/Sorghum | Cotton | 2024/08/22 | PV699146 |  |  |
| F29 | *F. proliferatum* | Moore | Corn | Cotton | 2024/05/01 | PV717848 |  |  |
| F13 | *F. clavum* | Hansford | Cotton | Wheat | 2024/05/01 | PV699143 |  |  |
| F26 | *F. clavum* | Randall | Corn/Sorghum | Cotton | 2024/05/01 | PV717846 |  |  |
| F27 | *F. clavum* | Hartley | Corn | Cotton | 2024/09/26 | PV717835 |  |  |
| F17 | *F. oxysporum* | Randall | Corn/Sorghum | Cotton | 2024/08/22 | PV699147 | PV731270 | PV699150 |
| F34 | *F. oxysporum* | Carson | Corn | Cotton | 2024/10/24 | PV717849 | PV731268 | PV699151 |
| F39 | *F. flocciferum* | Carson | Corn | Cotton | 2024/10/24 | PV717851 |  |  |
| F24 | *Fusarium sp.* | Sharman | Corn | Cotton | 2024/08/22 | PV699156 |  |  |
| F28 | *Fusarium sp.* | Hansford | Wheat | Cotton | 2024/05/01 | PV717847 |  |  |

The PCR products were detected using 1% agarose gel in 1x TAE buffer. The PCR products were cleaned using DNA Clean & Concentrator kit (Zymo Research) and sent to Eurofins Genomics (Eurofins Genomics LLC, Louisville, Kentucky) for sequencing with both forward and reverse primers. ABI sequence chromatograms were edited using SnapGene 8.0 (https://www.snapgene.com/) and exported as FASTA files that were aligned using MUSCLE alignment (Edgar 2004) in Mega11 (https://www.megasoftware.net/). Consensus sequences were obtained using BioEdit 7.7.1 (http://www.mbio.ncsu.edu/BioEdit/bioedit.html), which were then used to conduct BLASTn queries of NCBI GenBank (https://www.ncbi.nlm.nih.gov/genbank/) to obtain preliminary identifications.

Given the regional importance of *F. oxysporum*, two isolates identified as *F. oxysporum* were additionally amplified using RNA polymerase II second largest subunit (RPB2) (Liu et al. 1999) with the primer pair: 5f2 (GGGGWGAYCAGAAGAAGGC) × 7cr (CCCATRGCTTGYTTRCCCAT), using the PCR conditions described above, to further confirm its identity.

The two isolates identified as *F. oxysporum* in the present study were tested to determine whether they are FOV4 (and to determine whether they are genotype N, T, MT, and MiT) using specific primer pairs developed by Bell et al. (2019), following the protocol described by Wagner et al. (2022). This PCR detection method uses a 500 bp primer set that is based on the Tfo1 transposon insertion in the N-acetyltransferase (NAT) gene to differentiate FOV4/7 (VCG0114) from other races or VCGs. The primers used to genotype (T, N, MT, MiT) are based on the presence (T) or absence (N) of the Tfo1, MULE (MT) or MITE (MiT) insertions in the phosphate permease (PHO) gene (Wagner et al. 2022). Specific primer pairs included: FovT-F (GGCCGATATTGTCGGTCGTA) x FovA-R (TCAACAGACCCTGCACTACG) for the detection of FOV4; FovP-F (GGCCGATATTGTCGGTCGTA) x FovP-R (CTCCAGTGCAGTGCTTGGTA) for the detection of FOV4 N genotype; FovP-F (GGCCGATATTGTCGGTCGTA) x FovT-R (ATCTGTCTTTCGTCGGCAAT) for the detection of FOV4 T or FOV4 MiT genotype; and FovP-F (GGCCGATATTGTCGGTCGTA) x FovM-R (CCGCCATATCCACTGAACA) for the detection of FOV4 MT genotype.

### Phylogenetic analyses

Nucleotide sequences generated in this study, along with reference sequences retrieved from the NCBI GenBank database, were aligned using MUSCLE alignment (Edgar 2004) in Mega11 (https://www.megasoftware.net/). Alignments were then trimmed avoiding missing data at either end of alignments. The aligned sequences were analysed with maximum likelihood bootstrapping (MLBS) using IQ-TREE (http://www.iqtree.org/) (Nguyen et al. 2015) to determine the best-fit model of molecular evolution. These models were based on the Bayesian information criterion (BIC) scores (Chernomor et al. 2016) using ModelFinder (Kalyaanamoorthy et al. 2017). Once the optimal models were determined, clade support was assessed via 50,000 ML-BS pseudoreplicates of the data using IQ-TREE.

To determine the phylogenetic placement of *Fusarium oxysporum* sequences generated in this study relative to known FOV races (Cianchetta et al. 2015; Wang et al. 2010) and closely related sequences, additional reference sequences representing recognized FOV races and top BLASTn hits from the NCBI GenBank database were retrieved and phylogenetic analysis was conducted as described above. DNA sequences generated in the present study were deposited in NCBI (Table 1).

### Pathogenicity trials

A single trial was conducted to test the pathogenicity and severity of seven isolates, each representing a different *Fusarium* species, on corn and cotton. These included *F. incarnatum, F. equiseti, F. solani, F. verticillioides, F. proliferatum, F. clavum, and F. flocciferum.* Given its regional importance, *F. oxysporum* was not included in the inoculated pathogenicity trials due to regional concerns in relation to potentially working with FOV and the potential risk of its spread into these recently expanded cotton-growing areas.

### Pathogenicity of *Fusarium* species on cotton Biological material and experimental design

The trial was conducted under greenhouse conditions at Texas A&M AgriLife Research, Bushland, Texas at ambient temperatures (25 ± 5°C). A commonly cultivated variety in the Texas High Plains, FiberMax 868AXTP, with fair resistance to Fusarium wilt was used in this study (BASF 2024). Disease-free seeds of this variety were sourced from a previous trial: seeds were de-linted but received no chemical treatment prior to planting. Briefly, 10-cm pots were filled with a modified media (1 bag of Berger’s OM1 potting mix, 0.085 m^3^ of organic growing mix, 2.2 kg of sand, and 3 kg of soil). A total of 2-3 disease-free seeds were sown in each pot. Upon germination, seedlings were thinned to one per pot. There were six replicate pots for each *Fusarium* species and a non-inoculated control (total 48 plants), arranged in a completely randomized block design. Throughout the trial period, seedlings were fertilized with water-soluble fertilizer ‘Miracle-Gro’ as needed.

### Inoculum preparation

The sorghum grain inoculation method was used to prepare the inoculum (Yadav et al. 2017). Disease-free sorghum grains were sourced from a previous year’s trial.

Briefly, sorghum grains were soaked overnight, and 50 g of soaked grains were placed into each 250 mL beaker and sealed with aluminium foil. The beakers were autoclaved at 121°C for 30 minutes on two consecutive days. Once cooled, the sorghum grains in each beaker were inoculated with five agar plugs (5 mm), which were cut from a 7-day-old colony of specific *Fusarium* isolate grown on PDA. The inoculated beakers were gently shaken to uniformly distribute the inoculum and then placed inside zip-lock plastic bags to avoid contamination. The inoculated beakers were incubated for 7 days at room temperature (23 ± 2°C) with a 12-hour photoperiod and shaken every 2–3 days to ensure even distribution of the inoculum.

### Inoculation method

Cotton seedlings were inoculated at 2–3 leaf stage. Seedlings were watered thoroughly in the evening prior to inoculation the following morning. To inoculate seedlings, four 5-cm deep holes were made vertically into the potting medium of each pot using a wooden rod (18 x 0.8 cm), approximately 2 to 4 cm from the crown of each plant. A total of 1.25 g inoculum was inserted into each hole, excluding the non-inoculated controls. The holes were then covered with potting medium, and plants were watered daily following inoculation.

### Disease assessment

Cotton seedlings were observed daily for the appearance of disease symptoms. Disease severity on aerial plant parts was recorded at 7, 14, 21, and 28 days post-inoculation using a 0–4 rating scale, where: 0 = healthy; 1 = cotyledons wilted only; 2 = ≤50% true leaves wilted; 3= >50% but ≤90% true leaves wilted; and 4 = all leaves wilted or plant death (Table S1).

All cotton seedlings were harvested after 30 days of inoculation. Roots were washed free of soil and disease severity was recorded using 0–5 rating scale, where 0 = no symptoms; 1 = <10% of roots discolored; 2 = 10–30% of roots discolored; 3= 31–50% of roots discolored; 4 = 51–80% of roots discolored; and 5 = 81–100% of roots discolored (Table S1).

At the end of the trial, the roots of all cotton seedlings were plated onto PDA (1000 mL RO water, 39 g potato dextrose agar, 1 g streptomycin) to confirm pathogenicity.

### Pathogenicity of *Fusarium* species on corn

#### Biological material, experimental design, inoculum preparation and inoculation method

A commonly cultivated drought tolerant, untreated corn hybrid, Blue River Organic 82-14P, was used in the pathogenicity trial.

The experimental design, inoculum preparation and application method, as well as cultural practices, were the same as those used in the cotton pathogenicity trial. That is, a single batch of inoculum was used to inoculate both cotton and corn seedlings at the same time and similar cultural practices were followed to ensure consistency.

Seedlings were inoculated at the three-leaf stage.

## Disease assessment

Corn seedlings were harvested after one month of inoculation. Roots were washed free of soil and crown and root lesions were rated using a 1–8 rating scale (Parikh et al. 2018), where: 1 = no visible symptoms, 0% infection, and normal development; and 8 = seedling barely emerged and could not grow, scanty root but no emergence, or seed is dead (no emergence and no roots); infection coverage of seedling is >95% (Table S1).

At the end of the trial, the roots of all corn seedlings were plated onto PDA (1000 mL RO water, 39 g potato dextrose agar, 1 g streptomycin) to confirm pathogenicity.

## Data Analysis

In the case of disease severity recorded on aerial parts of cotton, the area under the disease progress stairs (AUDPS) was calculated using the ‘audps()’ function from ‘agricolae’ R package (Mendiburu 2023), based on severity ratings recorded across all four disease assessments. The AUDPS was selected over the traditional area under the disease progress curve (AUDPC) because the latter underestimates the effect of disease severity recorded during first and last disease assessments (Simko and Piepho 2012).

Linear mixed models effect models (LMMs) were then fitted using the ‘lmer()’ function from the ‘lmerTest’ R package (Kuznetsova et al. 2022) to investigate differences in disease severity among *Fusarium* species. The AUDPS was included as the dependent variable, the predictor *Fusarium* species were included as a fixed effect, and different blocks were included as a random effect to account for variability across blocks. Estimated marginal means for each *Fusarium* species were calculated using the ‘emmeans()’ function from ‘emmeans’ R package (Lenth 2021). The estimated marginal means were compared using Tukey-adjusted pairwise comparisons, and groupings were assigned using the ‘cld()’ function from the ‘multcomp’ R package (Hothorn 2020). These groupings indicated significant differences at α = 0.05. Models diagnostics was performed using the ‘DHARMa’ R package, which uses a simulation-based approach via the function ‘simulateResiduals’ to create readily interpretable scaled residuals for models (Hartig 2020).

In the case of disease severity recorded on roots of cotton and corn, differences in disease severity were tested using a nonparametric Kruskal-Wallis test using ‘ggbetweenstats()’ function from the ‘ggstatplot’ package (Patil 2021). If significant differences were found (α = 0.05), pairwise differences were performed using a post hoc Dunn test.

## Results

### Diversity and distribution of *Fusarium* species in the Northern High Plains

A total of nine *Fusarium* species were identified: these included *F. incarnatum, F. equiseti, F. solani, F. verticillioides, F. proliferatum, F. clavum, F. oxysporum, F. flocciferum,* and *Fusarium* sp. (Table 1, Figure 1). The number of isolates recovered for each species and their counties of origin are given in Table 1. Colony morphology (e.g., mycelial growth patterns) on PDA varied among isolates belonging to the same species, particularly for *F. solani. Fusarium solani* was the most frequently isolated species (39%), followed by *F. equiseti* (22%), *F. proliferatum* and *F. clavum* (7% each), *F. incarnatum*, *F. verticillioides*, *F. oxysporum*, and *Fusarium* sp. (5% each), and *F. flocciferum* (2%) (Table 1).

**Fig. 1.**
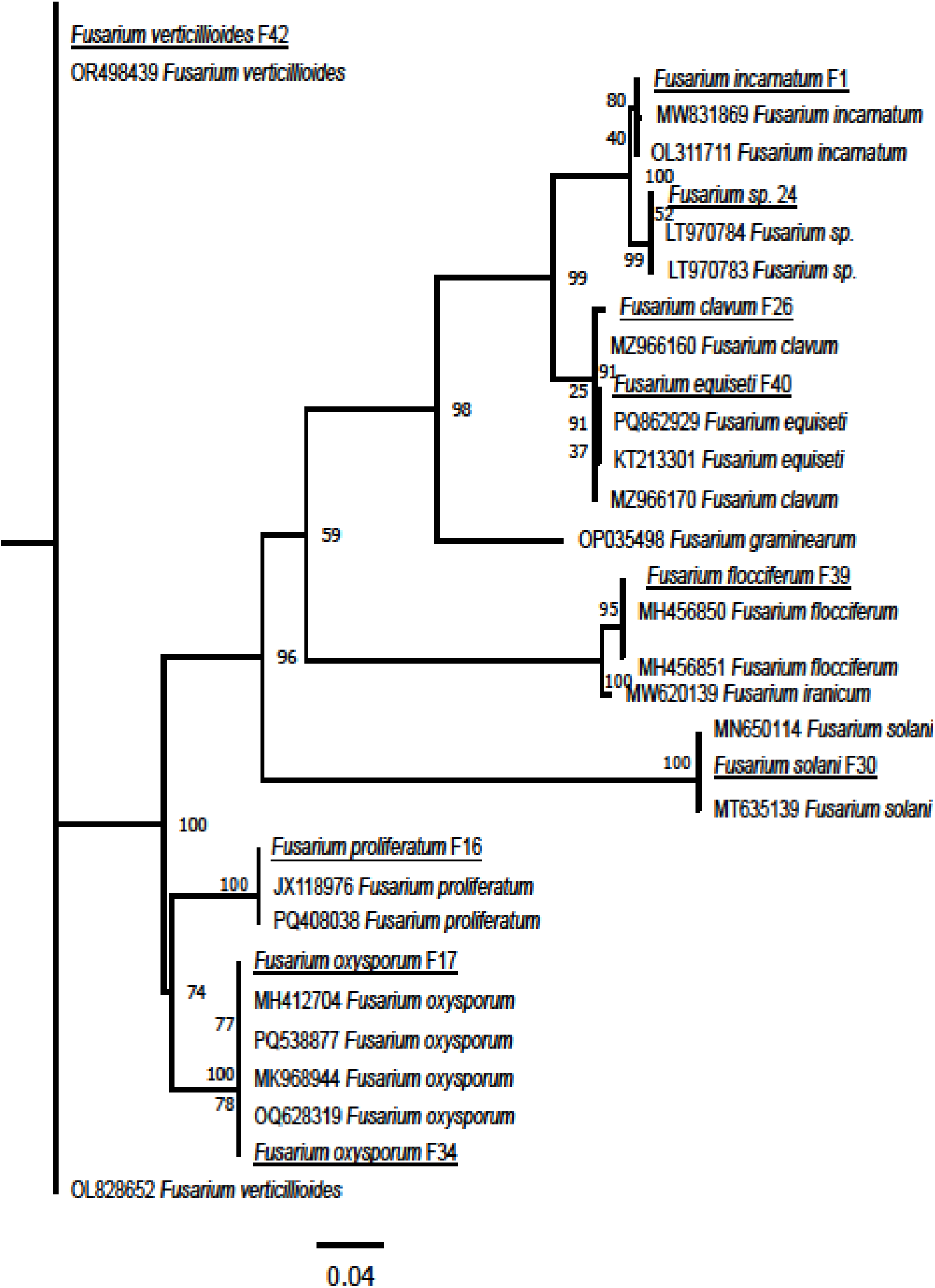
Phylogenetic comparison of *Fusarium* species isolated from cotton-corn rotation fields in the Texas High Plains (underlined) in relation to closely related reference species based on the translation elongation factor 1-α (tef-1α) gene region.

No PCR amplifications were achieved for two *F. oxysporum* isolates identified in this study using FOV4 specific primers. Further confirmation that the two *F. oxysporum* isolates were not FOV4 can be observed from the phylogenetic analysis of tef-1α gene region, which placed these two isolates within a clade containing reference sequences of *F. oxysporum* f. sp. *asparagi* and *F. oxysporum* f. sp. *lactucae* (Figure S1).

Although no single county harbored all species isolated in this study, five species were recovered from Sherman County: *F. incarnatum*, *F. equiseti*, *F. solani*, *F. verticillioides*, and *Fusarium* sp. (Table 1). This was followed by Moore County (*F. incarnatum*, *F. equiseti*, *F. solani*, and *F. proliferatum*) and Carson County (*F. equiseti*, *F. solani*, *F. oxysporum*, and *F. flocciferum*), each with four species isolated (Table 1).

Two species were restricted to a single county. That is, *F. verticillioides* was isolated only from Sherman County, while *F. flocciferum* was isolated from Carson County only (Table 1).

### Pathogenicity of *Fusarium* species on cotton

All *Fusarium* isolates tested were pathogenic on cotton; however, disease severity was largely restricted to root discoloration (Figure 2). No disease was recorded on non-inoculated control plants. *Fusarium* isolates were re-isolated from all inoculated roots, both symptomatic and asymptomatic, plated on PDA (i.e., 100% re-isolation rate).

**Fig. 2.**
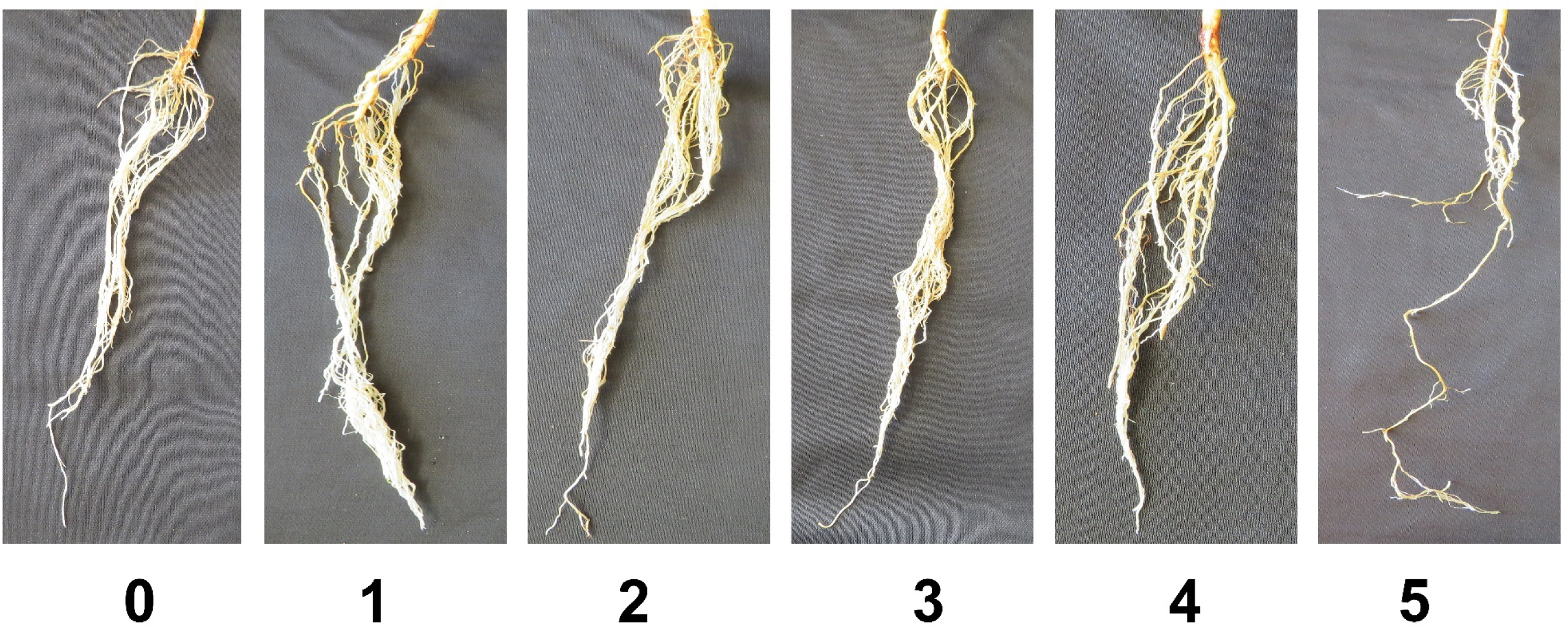
Damage on roots of cotton seedlings inoculated with *Fusarium* species. Whole root systems were rated visually using a 0–5 rating scale. Left to right: 0 - no symptoms; 1 - <10% of roots discolored; 2 - 10–30% of roots discolored; 3 - 31–50% of roots discolored; 4 - 51–80% of roots discolored; and 5 - 81–100% of roots discolored.

For the aerial parts, the highest disease severity (expressed as AUDPS) was caused by *F. incarnatum* (31.4), followed by *F. solani* (12.5), while the lowest disease severity was associated with *F. verticillioides* (7.1) (Figure 3A). Only *F. incarnatum* caused significantly higher disease severity relative to non-inoculated control (Figure 3B, *P*=0.0137, EMMeans pairwise comparisons). Disease severity associated other *Fusarium* species did not differ significantly from the non-inoculated control plants or from each other (Figure 3B, *P*>0.05, EMMeans pairwise comparisons). DHARMa diagnostics showed that the data met the model assumptions.

**Fig. 3.**
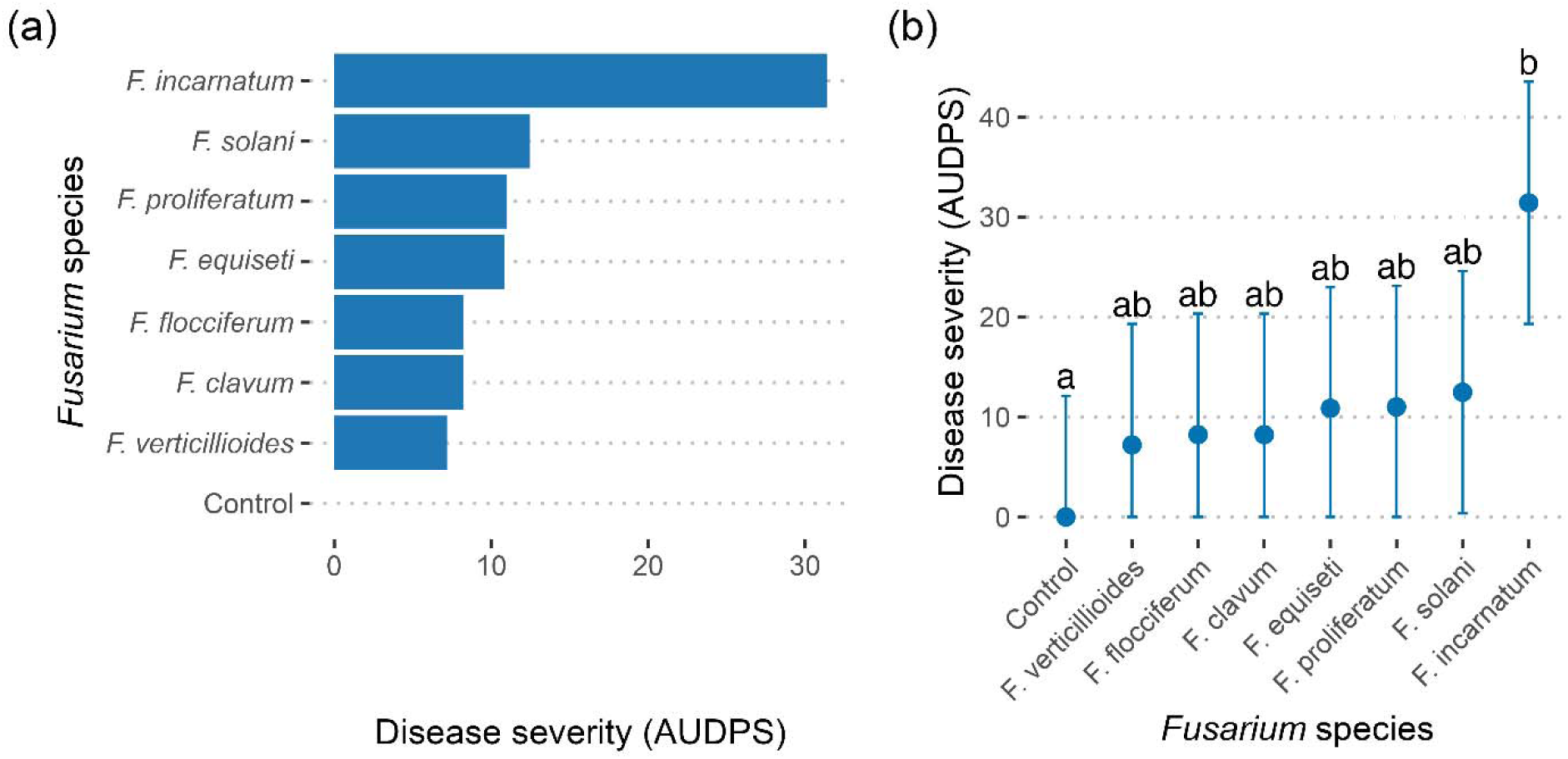
Comparison of the area under the disease progress stairs (AUDPS) based on severity ratings recorded on aerial parts of cotton seedlings across all four disease assessments: (A) AUDPS for each *Fusarium* species; (B) a linear mixed effect model fit showing estimated marginal means using the emmeans R package. Different letters (a, b) indicate significant differences among treatments at *P*≤0.05.

For roots, the Kruskal-Wallis test indicated that there was a significant difference in disease severity among groups (Figure 4A, χ^2^ = 17.58, df = 5, *P* = 0.01). The highest disease severity, expressed as median rank, was caused by *F. proliferatum* and *F. clavum* (2 each) (Figure 4A)*. Fusarium flocciferum*, *F. proliferatum,* and *F. clavum* caused significantly higher disease severity relative to non-inoculated control (Figure 4A, *P=*0.01, *P=*0.01, *P=*0.02, respectively). However, no significant differences in disease severity were recorded among different *Fusarium* species (Figure 4A, *P*>0.05).

**Fig. 4.**
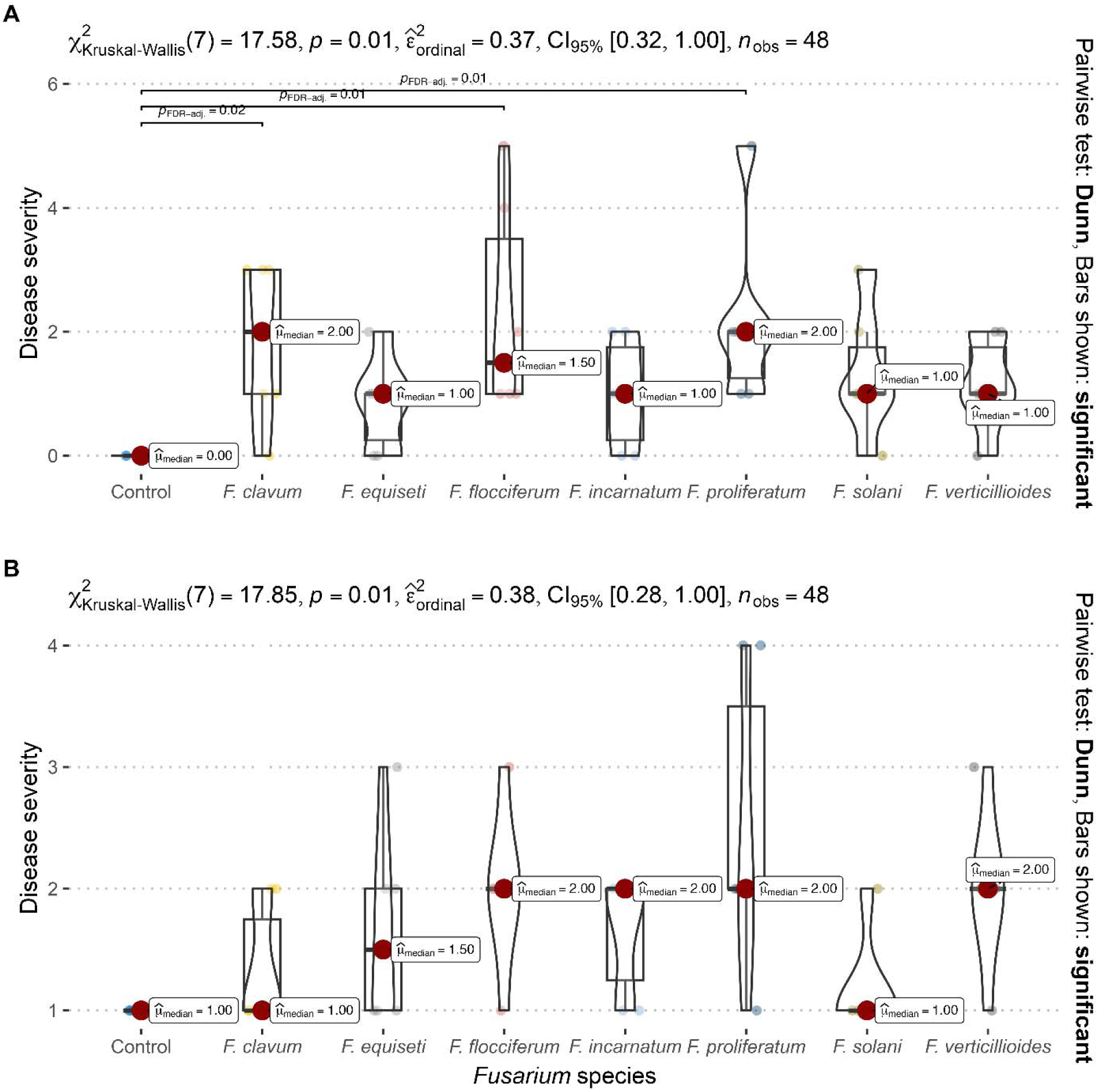
Comparison of disease severity of different *Fusarium* species on roots of (A) cotton and (B) corn. Differences in disease severity were compared using a non-parametric Kruskal–Wallis test.

### Pathogenicity of *Fusarium* species on corn

All *Fusarium* isolates tested were pathogenic on corn, causing stalk rot, root necrosis and brown to black discolorations; however, disease severity did not exceed 20–40%. The highest disease severity, expressed as median rank, was caused by *F. proliferatum, F.* flocciferum, *F. incarnatum* and *F. verticillioides* (2 each) (Figure 4B). No disease was recorded on non-inoculated control plants. Like cotton, *Fusarium* species were re-isolated from all inoculated corn plants, regardless of whether they exhibited symptoms. Differences in disease severity were not statistically significant among *Fusarium* species or relative to non-inoculated control plants (Figure 4B, *P*>0.05).

## Discussion

To the best of our knowledge, this study is the first to conduct a comprehensive survey of the recently expanded cotton-growing areas in the Northern High Plains of Texas and provided valuable insights into the diversity, distribution, and pathogenicity of *Fusarium* species associated with cotton-corn rotation fields. A total of nine *Fusarium* species were identified, with *F. solani* being the most frequently isolated and *F. flocciferum* the least. Sherman County harboured the highest number of *Fusarium* species. All tested species were pathogenic on cotton and corn during the pathogenicity assays conducted under greenhouse conditions.

The high frequency of *F. solani* and *F. equiseti* isolation in the present study aligns with prior findings from Arizona and Alabama, where they were the most frequently isolated species from cotton roots and rhizosphere soil (Mao and Hu 2019; Palmateer et al. 2002). Likewise, *F. solani* was reported as the most predominant species, isolated from 97% of sorghum and corn fields across central and southeastern United States (Leslie et al. 1990). Likewise, the least frequent isolation of *F. flocciferum* in the present study is consistent with the findings from previous studies conducted in France (Nguyen et al. 2024) and Belgium (Scauflaire et al. 2011), where it was among the least frequently isolated species in corn agroecosystems. To date, there are no reports of *F. flocciferum* as a widely distributed pathogen in the U.S. Cotton Belt. These findings suggest that *F. solani* and *F. equiseti* are habitat generalists, being widely distributed across the U.S. Cotton Belt, whereas *F. flocciferum* is mostly a habitat specialist with a narrow distribution range.

The observation that colony morphology (e.g., mycelial growth patterns) on PDA varied among isolates of the same species, particularly *F. solani,* is consist with finding from a previous study (Mao and Hu 2019) and suggests that morphological characteristics alone are insufficient for species identification, emphasizing the need for sequence-based identification.

The variation in *Fusarium* species diversity observed across counties in this study is likely caused by a combination of factors, including environmental conditions, anthropogenic disturbance, variation in the number of fields sampled across counties, soil types, cultural practices (e.g., crop rotation and tillage), and predisposing factors such as the presence of nematodes or flooding events (Mai and Abawi 1987; Steinkellner and Langer 2004). Differences in cotton acreage among counties may have also contributed to variation in species diversity across counties. Specifically, the highest species diversity in Sherman, Moore, and Carson counties can partly be attributed to the substantial increase in cotton acreage between 2013 and 2018, during which production grew by 301% to over 250,000 hectares (Ledbetter 2020; Marek and Bordovsky 2006).

The significantly higher disease severity observed on cotton inoculated with *F. flocciferum*, *F. proliferatum*, and *F. clavum*, and *F. solani* (relative to non-inoculated control plants), suggests that these species may pose a potential threat to cotton and corn production in the Northern High Plains. Among these, *F. proliferatum* and *F. solani* have been previously identified as the most virulent species on cotton (Abdel-Latif et al. 2007; Palmateer et al. 2002). *Fusarium proliferatum* has also been reported as a pathogen of cotton in New Mexico (Zhu et al. 2019). Disease severity on corn did not differ significantly between inoculated and control plants or among *Fusarium* species; however, *F. proliferatum* and *F. verticillioides* have been described as the most virulent species on corn in previous studies (Okello et al. 2019; Wang et al. 2021). The pathogenicity of *Fusarium* species on both cotton and corn suggests that they can overwinter on the stubble of either crop and serve as a primary source of inoculum in the next spring.

The observation of overall low disease severity on both cotton and corn may partly be attributed to the application of inoculum after seedling emergence for both cotton and corn, rather than the use of a root-dip inoculation method. Although the root-dip method can result in increased disease severity, it does not closely mimic natural infection under field conditions and may introduce stress during transplantation, potentially predisposing seedlings to infection (Diaz et al. 2021; Schoeneweiss 1975). Our findings suggest that the *Fusarium* species isolated in the present study may behave opportunistically, with plant stress playing an important role in increasing disease incidence under field conditions. This further implies that higher disease severity may be obtained in pathogenicity screening assays if seeds are planted directly into inoculated soil, rather than applying inoculum after seedling emergence.

The 100% re-isolation rate of *Fusarium* species from both symptomatic and asymptomatic roots following pathogenicity trials in the present study supports the previous findings that systemic colonisation can occur in the absence of visible disease symptoms (Leslie et al. 1990). This implies that the pathogenicity of *Fusarium* species in the field may need to be interpreted with caution, as *Fusarium* species may be living in plants as endophytes or secondary invaders (Leslie et al. 1990). In the case of plant tissues, this further underscores the importance of sampling both symptomatic and asymptomatic plant tissues for accurately determining the diversity and distribution of *Fusarium* species (Leslie et al. 1990). The present study likely provided an accurate estimate of *Fusarium* species diversity and distribution because isolates were recovered from soil samples, which minimized the bias associated with symptom-based tissue sampling and isolation. Furthermore, a previous study did not find any significant differences in the number of *Fusarium* species isolated from cotton roots and rhizosphere soil (Mao and Hu 2019), suggesting that soil-based isolation approaches can reliably capture species diversity.

This study provided baseline *Fusarium* species in the Northern High Plains of Texas. The detection of both common (e.g., *F. solani* and *F. equiseti*) and less frequently reported species (e.g., *F. clavum* and *F. flocciferum*) highlights the diversity of *Fusarium* species in this region. The non-detection of the highly aggressive FOV4, using FOV4 specific primers, in present study suggests that the pathogen may not yet have been introduced into the Northern High Plains. Additionally, the phylogenetic clustering of the two *F. oxysporum* isolates recovered in the present study with reference sequences of *F. oxysporum* f. sp. *asparagi* and *F. oxysporum* f. sp. *lactucae* suggest that they unlikely belong to the forma specialis ‘vasinfectum’. Therefore, there is a probability that *F. oxysporum* isolates recovered in this study may have not been pathogenic on cotton. However, given the known presence of FOV4 in nearby regions such as El Paso, Hudspeth, and New Mexico, continued surveillance remains essential.

Crop rotation is unlikely to be effective since *Fusarium* species are cross-host pathogenic and the region is primarily a rangeland where mainly susceptible crops such as cotton, corn and wheat are grown for fodder. For species with broad niches, containment and treatment (such as soil solarization) would potentially be required over vast areas and therefore be impractical. Local containment may still be an option for species with restricted niches. However, the use of resistant or tolerant cultivars should be prioritized as a primary disease management strategy.

Since *Fusarium* species diversity can vary with the time of the year (Fulcher et al. 2021), future studies should aim to collect samples across multiple seasons. Additionally, including counties not sampled in the present study will contribute to a more comprehensive understanding of *Fusarium* species diversity and distribution across the Northern High Plains of Texas.

## Acknowledgements

We would like to acknowledge Dennis Coker, Marcel Fischbacher, JD Ragland, Kristy Slough, Josie Strnad, and Laura Taylor for assistance in field selection, and Kristen Tatro for greenhouse technical support. The work was supported by funding from Cotton Inc. and Texas A&M AgriLife Research.

## Author-Recommended Internet Resources

https://github.com/IhsanKhaliq/HighPlainsFusa

---

**Fig. S1.**
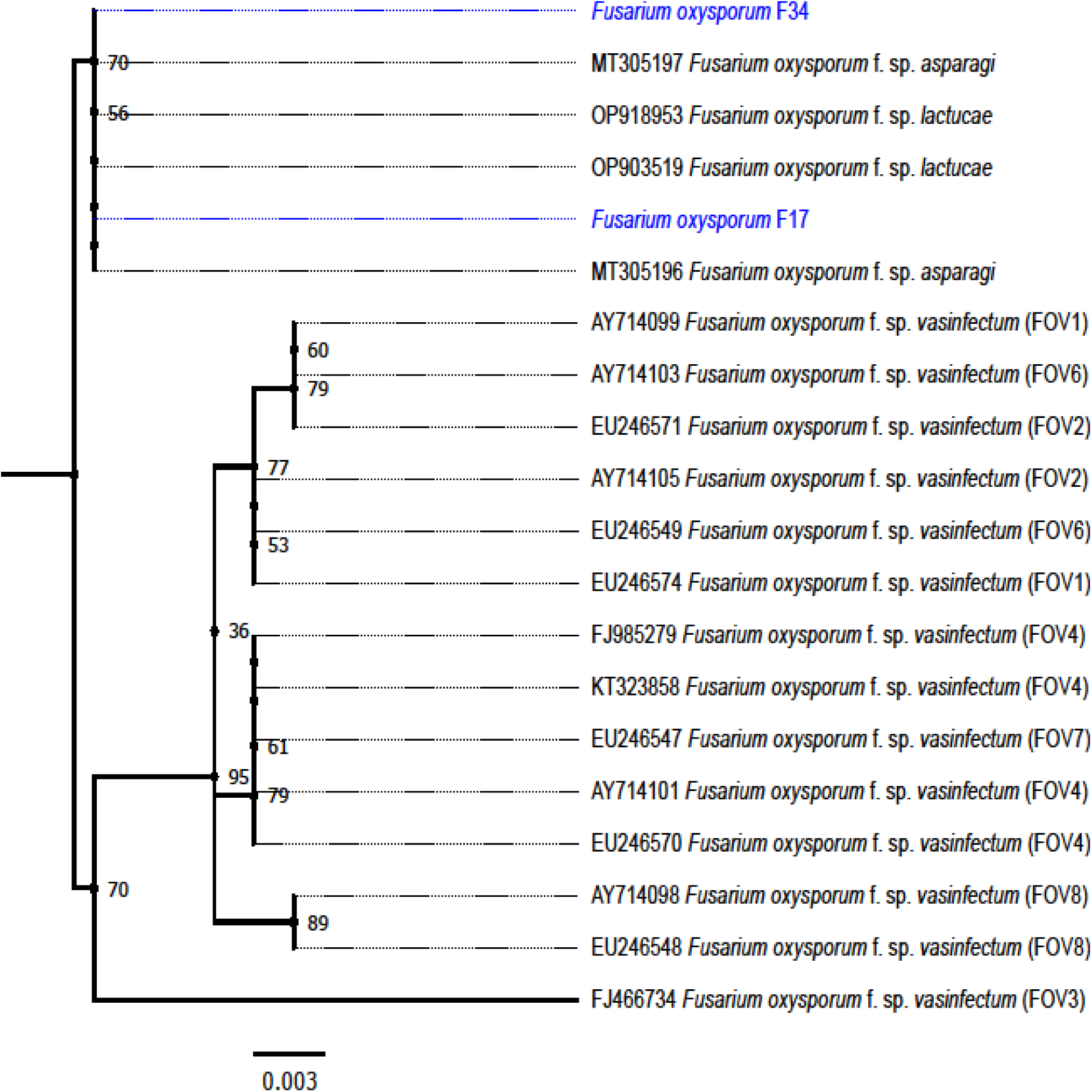
*Phylogenetic* relationships of *Fusarium oxysporum* isolates collected in this study (highlighted in blue) compared with reference isolates of *F. oxysporum* f. sp. *vasinfectum* races 1, 2, 3, 4, 6, and 8 (FOV1 to FOV8) and other closely related species, based on sequences of the translation elongation factor 1-α (tef1-α) gene region.

**Table S1.**
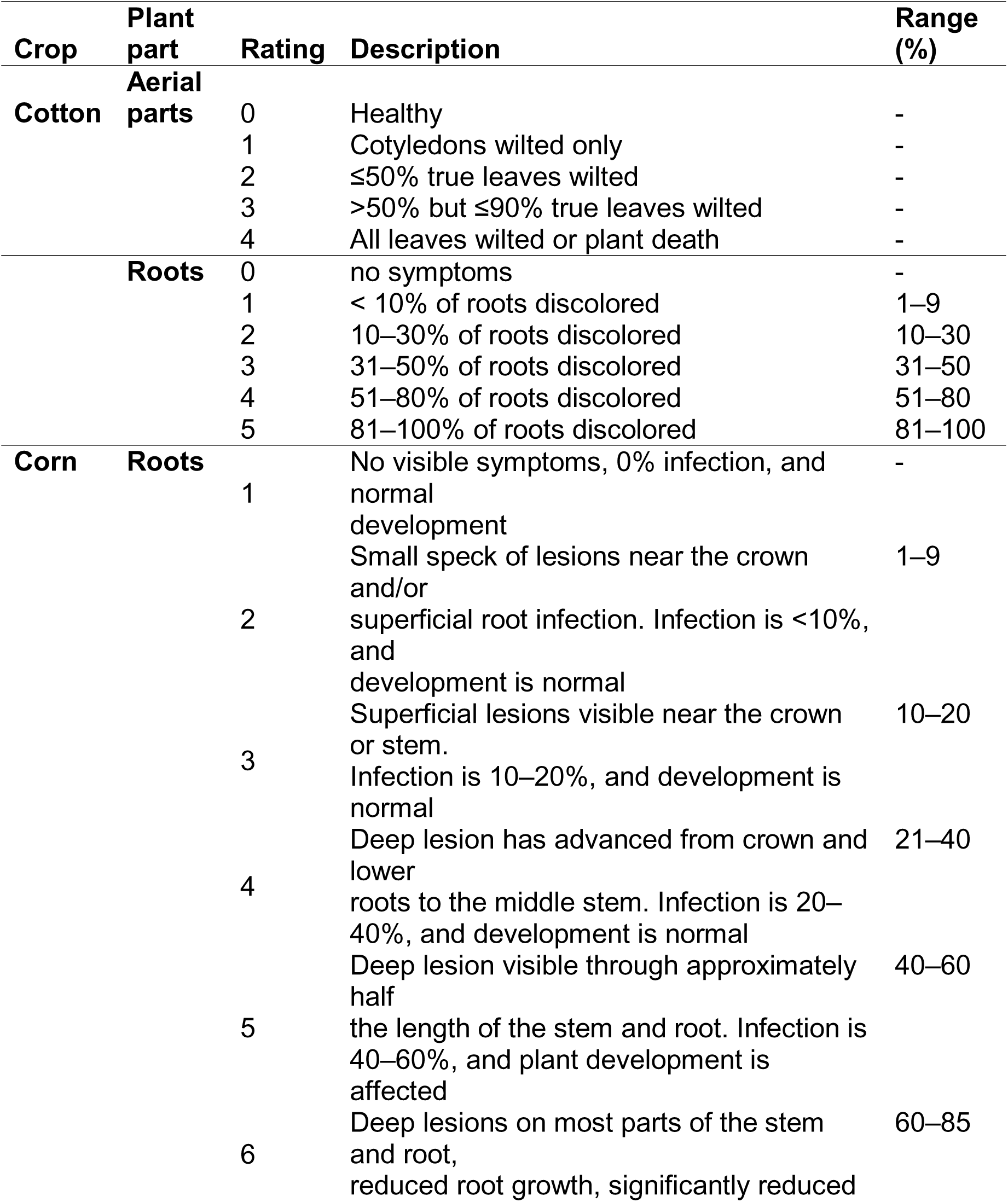
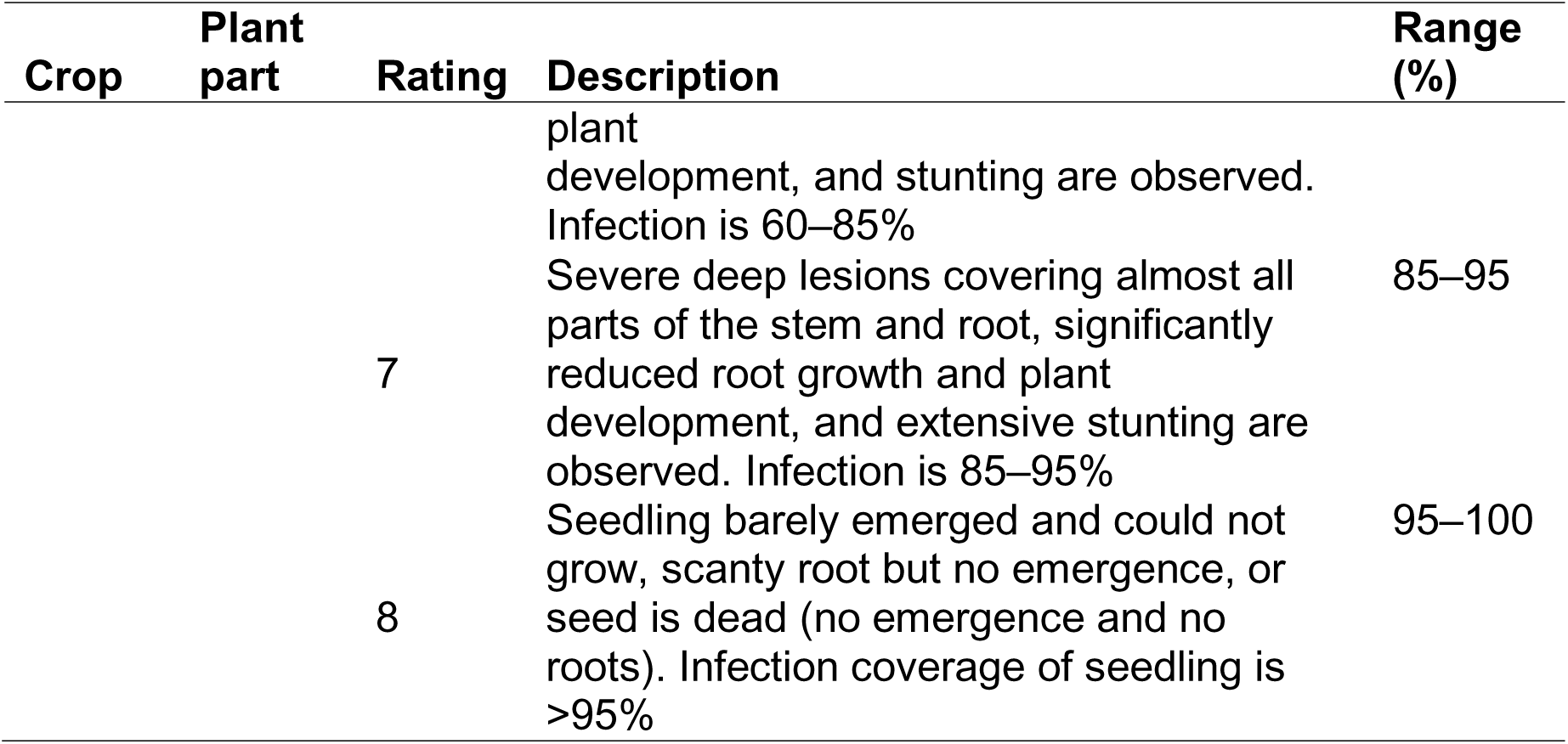
Disease rating scales used for assessing disease severity of *Fusarium* species in cotton and corn. For cotton, disease severity was recorded on both aerial parts and roots, whereas for corn, it was recorded on roots only.

